# Combining machine learning and a universal acoustic feature-set yields efficient automated monitoring of ecosystems

**DOI:** 10.1101/865980

**Authors:** Sarab S. Sethi, Nick S. Jones, Ben D. Fulcher, Lorenzo Picinali, Dena J. Clink, Holger Klinck, C. David L. Orme, Peter H. Wrege, Robert M. Ewers

## Abstract

Natural habitats are being impacted by human pressures at an alarming rate. Monitoring these ecosystem-level changes often requires labour-intensive surveys that are unable to detect rapid or unanticipated environmental changes. Here we developed a generalisable, data-driven solution to this challenge using eco-acoustic data. We exploited a convolutional neural network to embed ecosystem soundscapes from a wide variety of biomes into a common acoustic space. In both supervised and unsupervised modes, this allowed us to accurately quantify variation in habitat quality across space and in biodiversity through time. On the scale of seconds, we learned a typical soundscape model that allowed automatic identification of anomalous sounds in playback experiments, paving the way for real-time detection of irregular environmental behaviour including illegal activity. Our highly generalisable approach, and the common set of features, will enable scientists to unlock previously hidden insights from eco-acoustic data and offers promise as a backbone technology for global collaborative autonomous ecosystem monitoring efforts.

## Introduction

With advances in sensor technology and wireless networks, automated passive monitoring is growing in fields such as healthcare^1^, construction^2^, surveillance^3^ and manufacturing^4^ as a scalable route to gain continuous insights into the behaviour of complex systems. A particularly salient example of this is in ecology, where due to accelerating global change^5^, we urgently need to track changes in ecosystem health, accurately and in real-time, in order to respond to ecosystem threats^6,7^. Traditional ecological field survey methods are poorly suited to this challenge: they tend to be slow, labour intensive, narrowly focused and are often susceptible to observer bias^8^. Using automated monitoring to provide scalable, rapid, and consistent data on ecosystem health^9,10^ seems an ideal solution, yet progress in implementing such solutions has been slow. Existing automated systems tend to retain a narrow biotic or temporal focus and do not transfer well to novel ecosystems or threats^11,12^.

We present an innovative framework for automated ecosystem monitoring using eco-acoustic data (**Fig. 1**). We used a pre-trained general-purpose audio classification convolutional neural net (CNN)^13,14^ to generate acoustic features, and discovered that these are remarkably powerful ecological indicators that are highly descriptive across spatial, temporal and ecological scales. We were able to discern acoustic differences among biomes, detect spatial variation in habitat quality, and track temporal biodiversity dynamics through days and seasons with accuracies far surpassing that possible using existing methods. We extended this approach to demonstrate efficient exploration of large monitoring datasets, and the unsupervised real-time detection of anomalous environmental behaviour, including hallmarks of illegal activity.

**Figure 1:**
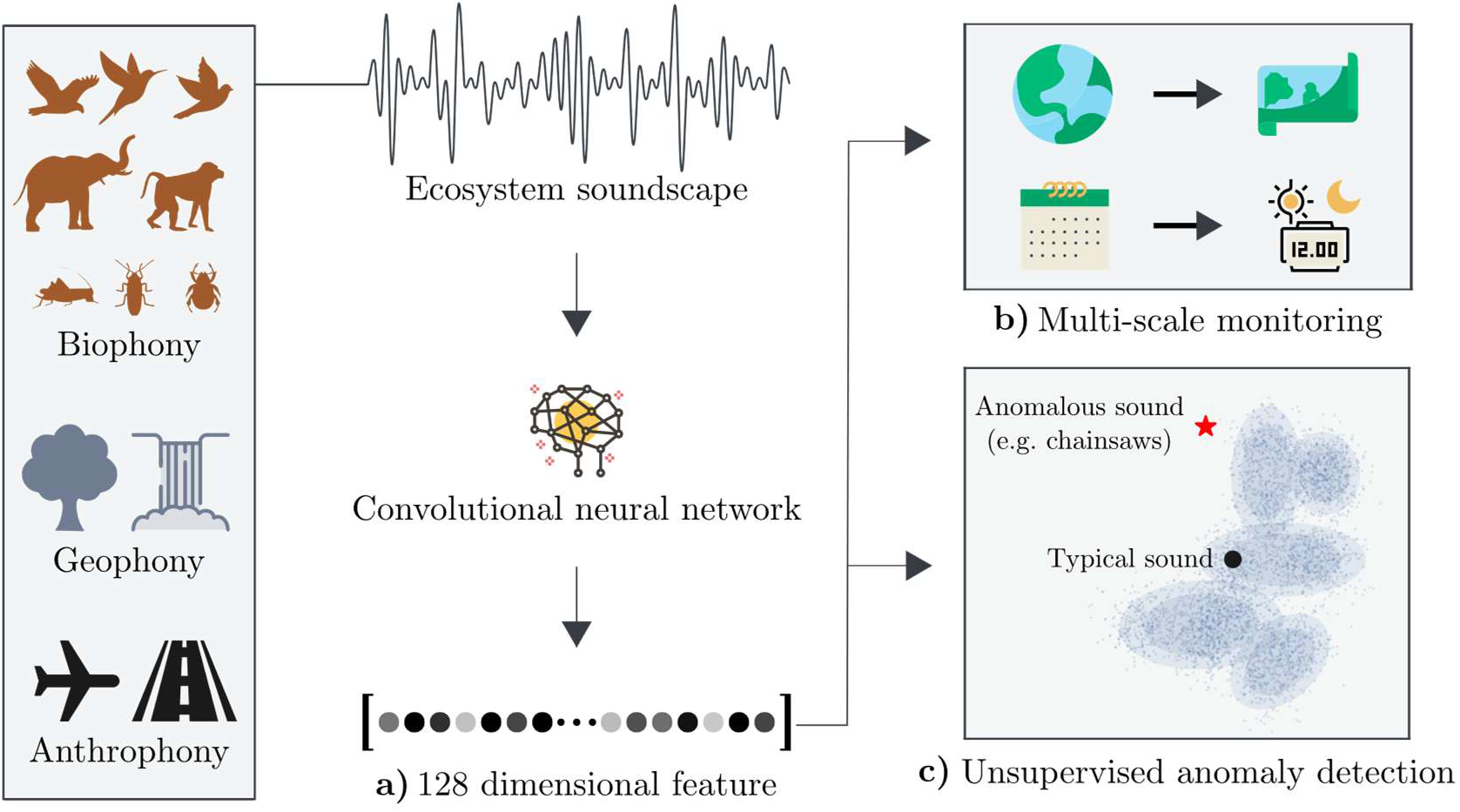
A common framework for monitoring ecosystems autonomously using soundscape data.

**(a)** We embed eco-acoustic data in a common high-dimensional feature space using a convolutional neural network (CNN). Remarkably, this common embedding means that we can both **(b)** draw out ecological insights into ecosystem health across multiple temporal and spatial scales, and **(c)** effectively identify anomalous sounds in an unsupervised manner.

Our approach avoids two pitfalls of previous algorithmic assessments of eco-acoustic data^15^. First, we do not require supervised machine-learning techniques to detect^16,17^ or identify^18,19^ target threats or species. Supervised methods use annotated training datasets to describe target audio exemplars. This approach can yield high accuracy^20^, but is narrowly focused on the training datasets used, can be subverted (e.g. in the case of illegal activity detection^21^), requires investment in laborious data annotation, and frequently transfers poorly from one setting to another^22^.

Second, we do not depend on specific hand-crafted eco-acoustic indices. Such indices share our approach of aggregating information across a whole audio sample^23^ – a soundscape – but differ in their approach of identifying a small number of specific features (e.g. entropy of the audio waveform^24^) rather than a machine-learned, general acoustic fingerprint. Again, these indices can predict key ecological indicators in local contexts^25–27^, but they often fail to discriminate even large ecological gradients^28,29^, and behave unpredictably when transferred to new environments^30^.

A lack of transferability is characteristic of approaches that use site-specific calibration or training, which confers high local accuracy at the cost of generality. Lack of generalisability is a critical failure for monitoring applications, where rapid deployment is essential and the nature of both threats and responses cannot always be known in advance. Threats may be acute, such as logging or hunting^31^, or chronic, such as the invasion of a new species^32^ or climate change^33^, and may drive unpredictable ecological responses^34^. The remarkable efficacy of our feature-set means we discover a general solution to these complex methodological challenges; it is highly descriptive across spatial and temporal scales and is capable of reliably detecting anomalous events and behaviour across complex ecosystems.

## A common feature embedding yields multi-scale ecological insight

We collected global acoustic data from the following ecosystems: protected temperate broadleaf forests in both Ithaca, USA and Abel Tasman National Park, New Zealand; protected lowland rainforests in Sulawesi, Indonesia; protected and logged lowland rainforest in and surrounding Nouabalé-Ndoki National Park, Republic of Congo; and lowland rainforests across a gradient of habitat degradation in Sabah, Malaysia. In total we analysed over 2750 hours of audio, collected using a variety of devices including AudioMoths^35^, Tascam recorders, Cornell Lab Swifts, and custom set-ups using commercial microphones (Methods). We then embedded each 0.96 second sample of eco-acoustic data in a 128-dimensional feature space using a CNN pre-trained on Google’s AudioSet dataset^13,14^.

AudioSet is a collection of human-labelled sound clips, organised in an expanding ontology of audio events, which contains over two million short audio samples drawn from a wide range of sources appearing on YouTube. Although a small amount of eco-acoustic data is present, the vast majority of audio clips are unrelated to natural soundscapes^13^, with the largest classes consisting of music, human speech and machine noise. No ecological acoustic datasets provide labelled data on a similar magnitude to AudioSet, and when detecting ‘unknown unknowns’ it is in fact desirable to have a feature space that is able to efficiently capture characteristics of non-soundscape specific audio. The resulting acoustic features are therefore both very general and of high resolution, placing each audio sample in high-dimensional feature space that is unlikely to show ecosystem specific bias.

We investigated whether this embedding revealed expected ecological, spatial and temporal structure across our eco-acoustic datasets. Short audio samples are highly stochastic, so we first averaged acoustic features over five consecutive minutes. We were able to clearly differentiate eco-acoustic data from different ecosystems (**Fig. 2a**). Furthermore, samples from the same location clustered strongly, even when different recording techniques and equipment were used, and audio samples from similar ecosystems were more closely located in audio feature space (Fig. S1). Within sampling locations, the acoustic features clearly captured ecological structure appropriate to the spatial and temporal scale of recordings. Data recorded across a gradient of logging disturbance in Sabah^36^ reflected independent assessment of habitat quality based on the quantity of above ground biomass (AGB) (**Fig. 2b**), except for sites near rivers where the background sound of water dominated the audio. Monthly recordings from Ithaca captured eco-acoustic trajectories describing consistent seasonal changes in community composition driven by migratory fluxes of birds (**Fig. 2c**). Similarly, daily recordings in Sabah strongly discriminated between the dawn and dusk choruses in the tropical rainforest of Malaysia, with large discontinuities at 05:00 and 17:00 hours, respectively, that reflected diurnal turnover in the identity of vocalising species (**Fig. 2d**). The same acoustic features also revealed diurnal patterns in data from the four other biomes used in this study (Fig. S2). These results show that we are able to capture complex hierarchical structure in ecosystem dynamics using a common eco-acoustic embedding, with no modification required when moving across spatial and temporal scales.

**Figure 2:**
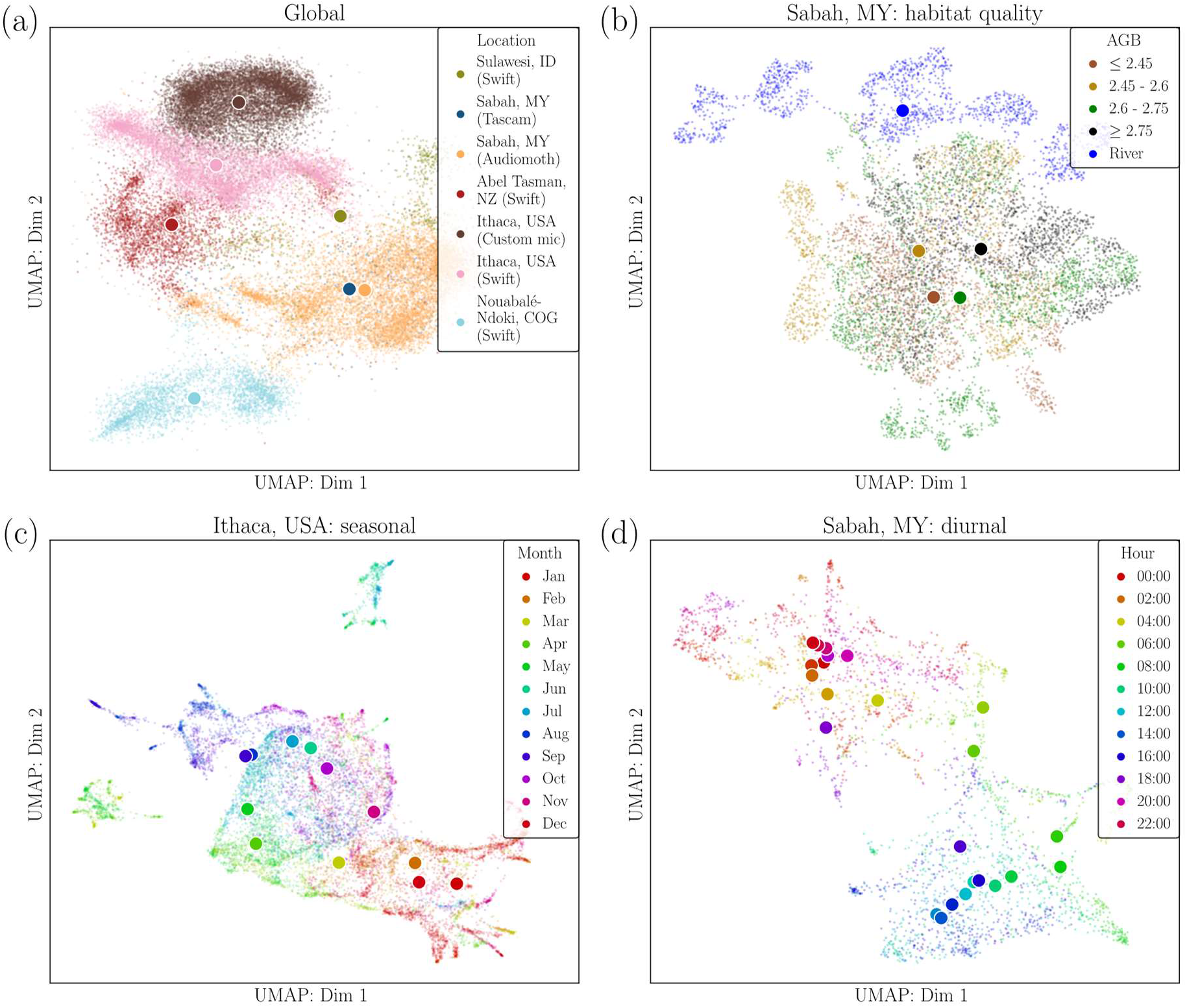
Embedding eco-acoustic data in a common, highly descriptive feature space yields ecological insight across spatial and temporal scales.

**(a)** Seven eco-acoustic datasets from five ecosystems around the world are embedded in the same acoustic feature space, in which different biomes are distinguished. Features were robust to different recording technologies used in Sabah (Tascam, Audiomoth) and Ithaca (Swift, custom microphone, Methods). **(b)** Tropical forest areas in Sabah that differ in habitat quality (measured by above ground biomass, log10(t.ha^−1^)) cluster using the same acoustic feature space. **(c)** Three years of soundscape data from a temperate forest in Ithaca reveals a clear seasonal cycle. **(d)** One month of acoustic data from a logged tropical forest site in Sabah shows a repeating diurnal pattern. In all panels, uniform manifold learning technique (UMAP)^37^ was used to visualise a 2D embedding from the full 128-dimensional acoustic feature space, and centroids of classes are denoted by larger points.

While unsupervised approaches can thus clearly be used to anatomize ecosystem data in our feature space, a core aim of autonomous monitoring systems is to directly predict ecosystem health, and to be able to do so longitudinally over long time periods. We showed that the same general acoustic features (derived from the pre-trained CNN) were well suited to this problem by performing a series of classification tasks. Classifications were performed using a random forest classifier in the full feature space, and we compared the performance (measured by F1 score^38^) with a feature space made up from five existing eco-acoustic indices (EAI) specifically designed to assess ecosystem health (Methods). Our approach provided markedly more accurate prediction of biodiversity and habitat quality metrics in both temperate (avian richness; CNN: 88% *versus* EAI: 59%; **Fig. 3a**) and tropical (AGB; CNN 94% *versus* EAI 62%; **Fig. 3b**) landscapes. Importantly, the prediction of avian richness did not require individual identification of species within the soundscape – a process only possible given vast amounts of manually labelled, species-specific data. General acoustic features also more accurately predicted temporal structure at both seasonal (months within temperate soundscapes; CNN 86% *versus* EAI 42%; **Fig. 3c**) and daily (hours within tropical soundscapes; CNN 68% *versus* EAI 31%; **Fig. 3d**) timescales. These results pave the way for automated eco-acoustic monitoring to detect environmental changes over long time scales. For example, the loss of tree biomass from logging over a period of months, annual shifts in the seasonal phenology of bird communities^39^, and the gradual increase of forest biomass through decades of forest recovery or restoration^40^, may all be accurately tracked through time using this analysis framework.

**Figure 3:**
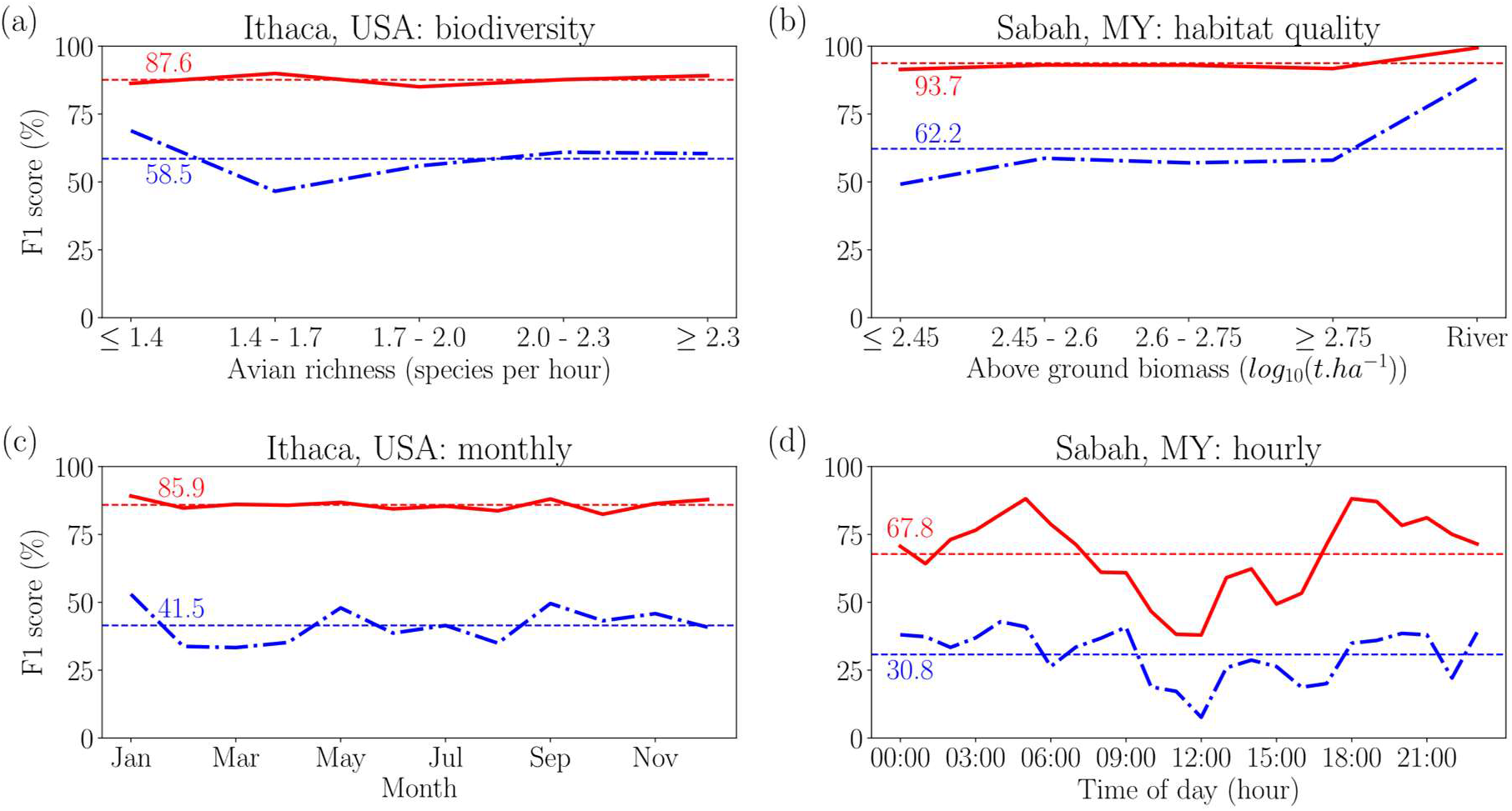
General acoustic features allow accurate classification of the degree of ecosystem degradation and position in diurnal and seasonal cycles.

We performed a multi-class classification task using a 20% test set to assess the predictive power of the general acoustic features on a range of spatial and temporal sales of eco-acoustic data. For each task we measured the F1 score for each of the classes, and compared the results using general acoustic features derived from a pre-trained CNN (red) to a baseline made up of standard eco-acoustic indices regularly used in eco-acoustics (blue, Methods). In **(a)** we were able to predict a measure of biodiversity (avian richness, species per hour) from a temperate forest site in Ithaca. **(b)** We are also able to predict habitat quality (as measured by above ground biomass, log10(t.ha^−1^)) across a landscape degradation gradient in tropical Malaysia with high accuracy, with the exception of sites near rivers. In **(c)** and **(d)** we show how temporal cyclicity on the scale of months and hours respectively can be predicted using the same acoustic feature-set.

## The common feature space allows effective unsupervised anomaly detection and eco-acoustic data summarisation

Given the huge volumes of audio data that are rapidly collected from autonomous monitoring networks, it is important to create automated summaries of these data that highlight the most typical or anomalous sounds at a given site – a task that is not possible given current approaches to eco-acoustic monitoring. In particular, the task of unsupervised anomaly detection is critical in real-time warning systems which need to automatically warn of unpredictable rapid changes to the environment, or illegal activities such as logging and poaching^31^. We solved both the problems of efficient data summarisation and unsupervised anomaly detection using a density estimation algorithm in our general acoustic feature space.

We developed a site-specific anomaly scoring algorithm using a Gaussian Mixture Model (GMM) fit to five full days of acoustic features from a given recording location. Here we used the original 0.96 second, 128-dimensional features, which best captured transient acoustic events (Methods). We then explored the most typical and anomalous sounds from a logged tropical forest in Sabah, Malaysia to demonstrate how this approach allows efficient exploration of large amounts of data (**Fig. 4a**, Methods). High-probability signals corresponded to distinct background noise profiles, driven primarily by insect and frog vocalisations which varied in composition throughout the day, but also included abiotic sounds such as rainfall. Low probability, or anomalous, sounds included sensor malfunctions, anthropogenic sounds such as speech, and biotic sounds such as species with distinctive calls that were heard rarely during the recording period (e.g. gibbon) or unusually loud species (e.g., a cicada immediately adjacent to the microphone) (**Fig. 4a**). Exploring the data in this way, we were able to acquire a high level, rounded summary of a 120 hour (432,000 s) period of acoustic monitoring, by listening to just 10 s of the most typical sounds and 12 s of anomalies (Supplementary Audio File 1).

**Figure 4:**
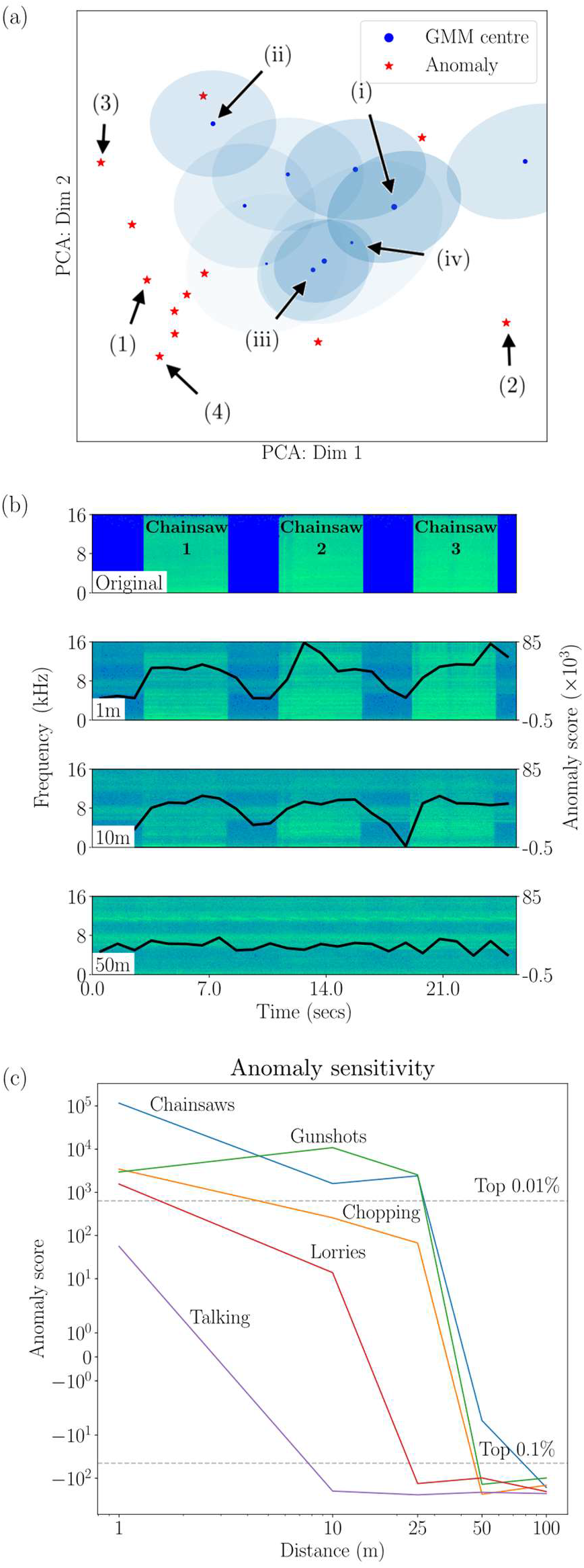
Density estimation in acoustic feature space allows unsupervised detection of anomalous sounds.

**(a)** A projection of the Gaussian Mixture Model (GMM) fit to five full days of data from one logged tropical forest site in Sabah, Malaysia. Principal component analysis was used to project the GMM centres and covariances from 128 to 2 dimensions for purposes of visualisation, and shaded areas correspond to two standard deviations from each GMM centre. Points close to the centres are typical background sounds, and thus given low anomaly scores (i, ii and iii = ambient noise at different times of day; iv = light rain in a largely silent forest; Supplementary Audio File 2). Conversely, very unusual sounds are in low density regions of acoustic space and are given high anomaly scores (1 = human talking; 2 = vocalising gibbon in background; 3 = sensor malfunction; 4 = loud insect near microphone, Supplementary Audio File 2). **(b)** We used playback experiments to test the sensitivity of the anomaly score to novel acoustic events, illustrated here by chainsaw sounds. Spectrograms are shown for audio recorded from the device when the anomalous audio file (original) was played from a speaker at a variety of distances. Blue and green represent time-frequency patches of low and high volume respectively, and overlaid in black is the anomaly score for each 0.96 s of audio. **(c)** We investigated the sensitivity of the algorithm to a variety of anomalous sounds typical of illegal activity (chainsaws, gunshots, chopping, lorries, talking). Anomaly scores were averaged across ten sites from a logged tropical forest landscape in Sabah and vary with distance of playback. Dotted lines show where averaged anomaly scores entered the top 0.1% and 0.01% respectively of all 449,280 0.96 s audio clips that were used to fit the probability density function.

Real-time detection of illegal human activities such as logging and poaching is a particularly pressing problem in protected areas^31^. One approach is to train supervised classifiers to search for sounds such as chainsaws^41^ or gunshots^42^, however not only do these require specific training datasets but they can easily go out of date or be subverted (e.g., by using a different gun^21^). We used calibrated playback experiments to test the efficacy of our unsupervised density estimation approach for detecting novel acoustic events without prior training. We used a speaker to play sounds including chainsaws, gunshots, chopping, lorries and speech at distances of 1, 10, 25, 50 and 100 m from an acoustic recorder within the ecosystem (**Fig. 4b**), and replicated this experiment across ten sites from the land degradation gradient in Sabah, Malaysia. All sounds were scored as strongly anomalous at 1 m, but differed in how the score declined with distance. Chainsaws and gunshots and, to a lesser extent, chopping all scored highly at distances of up to 25 m of forest from the recorder, but were not audible over background noise at greater distances (**Fig. 4c**). In contrast, lorries and speech were only reliably detected within about 10 metres of the recorder. Detection ranges in real-world settings will be larger as our playback experiments were unable to fully replicate the sound pressure levels of events such as gunshots (Methods). The same playback experiment also detected chainsaw and gunshot sounds in a temperate setting in Ithaca with no modification to the algorithm (Fig. S3), suggesting that this approach to automated anomaly detection is transferable among vastly differing ecosystems.

## The future of automated environmental monitoring

We have shown how state-of-the-art machine learning techniques can be used to draw out detailed information on the natural environment via its soundscape. Using a common feature-based embedding, derived from over 5000 hours of non-ecosystem audio data, we were able to monitor diverse ecosystems on a wide variety of spatial and temporal scales, and predict biological metrics of ecosystem health, with much higher accuracies than was previously possible from eco-acoustic data. Furthermore, we used the same approach to concisely summarise huge volumes of data, and identify anomalous events occurring in large datasets over long time periods, in an unsupervised manner. This novel approach offers a bridge from the mostly disappointing hand-crafted eco-acoustic indices and highly taxonomically specific detection-classification models to a truly generalisable approach to soundscape analysis. While here we have tackled the task of ecological monitoring, the same approach can easily be generalised to other fields employing acoustic analysis, for example in healthcare^1^, construction^2^, surveillance^3^ or manufacturing^4^. Pairing these new computational methods with networked acoustic recording platforms^43^ offers promise as a general framework on which to base larger efforts at standardised, autonomous system monitoring.

## Materials and methods

### Audio data collection

Audio data was collected from a wide variety of locations using different sampling protocols in this study.

In Sabah, Malaysia two datasets using different recording devices contained data across an ecological gradient encompassing primary forest, logged forest, cleared forest and oil palm sites^36^ collected between February 2018 and June 2019. In the Tascam dataset, audio was recorded as 20 minute sound files at 44.1 kHz using a Tascam DR-05 recorder mounted at chest height on a tripod, from 14 sites. One 20 minute file was recorded per hour, and a total of 27 hours 35 minutes was recorded. In the Audiomoth dataset, the Audiomoth device^35^ recorded continuously as five minute sound files at 16 kHz and the device was secured to a tree at chest height across 17 sites (14 overlapping with the Tascam dataset). A total of 748 hours of audio was recorded.

Two datasets were recorded from Sapsucker Woods, Ithaca, NY, USA using different methodologies. The first was from a single location, recorded almost continuously over three years, between January 2016 and December 2019 using a Gras-41AC precision microphone, and audio digitised through a Barix Instreamer ADC at 48 kHz. A total of 797 hours of audio was collected. The second contains 24 hours of audio from 13^th^ May 2017 and was recorded using 30 Swift acoustic recorders across an area of 220 acres. Audio was recorded as one hour files at 48 kHz and recorders were attached to trees at eye height. A total of 638 hours of audio was recorded.

In New Zealand, audio was recorded continuously using semi-autonomous recorders from the New Zealand Department of Conservation from 8^th^ to 20^th^ December 2016. Ten units were deployed in the Abel Tasman National park, with five on the mainland and five on Adele Island. Each sound file spanned 15 minutes and a sampling frequency of 32 kHz was used. Recorders were attached to trees at eye-height. A total of 240 hours of audio was recorded.

In Sulawesi, audio was recorded using Swift acoustic recorders with a standard condenser microphone in Tangkoko National Park, a protected lowland tropical forest area. Data was recorded continuously from four recording locations within the park during August 2018. Each recording spanned 40 minutes and a sampling frequency of 48 kHz was used. Recorders were set at 1m height from ground level. A total of 64 hours of data was recorded.

In the Republic of Congo, audio was recorded using Swift acoustic recorders with a standard condenser microphone from 10 sites in and surrounding Nouabalé-Ndoki National Park between December 2017 and July 2018. Sound files were 24 hours long, and a sampling frequency of 8 kHz was used. Habitat types spanned mixed forest and *Gilbertiodendron* spp. from within a protected area, areas within a six year old logging concession, and within active logging concessions. Recorders were set at 7-10 m from ground level, suspended below tree limbs. A total of 238 hours 20 minutes of audio was recorded.

### Acoustic feature embedding

Each 0.96 s chunk of eco-acoustic audio was first resampled to 16 kHz using a Kaiser window, and a log-scaled Mel-frequency spectrogram was generated (96 temporal frames, 64 frequency bands). Data recorded at below 16 kHz (Nouabalé-Ndoki, COG) was up-sampled to this frequency. This was then passed through a convolutional neural network (CNN) pre-trained on Google’s AudioSet dataset^13,14^, and a 128-dimensional embedding was computed as the output of the penultimate layer.

As a baseline comparison we created a similar embedding using a selection of standard soundscape metrics. These were Sueur’s α index^24^, temporal entropy^24^, spectral entropy^24^, Acoustic Diversity Index (ADI)^44^, and Acoustic Complexity Index (ACI)^45^. Each of the above features was computed over 1 s of audio and concatenated to create a five-dimensional feature vector. This is referred to as a compound index in standard eco-acoustic studies^25^.

For the multi-class classification problems, for prediction of biodiversity, and to create the visualisations in Fig. 2, we averaged both feature vectors over consecutive five-minute periods. The CNN we use from Hershey et. al.^14^ takes a Mel-scaled spectrogram of 0.96 s duration at a Nyquist frequency of 8 kHz as an input. Insects and bats in particular produce sounds reaching well into the ultrasonics^23^ which contain important ecological information but will be missed by this embedding. Similarly, the features may be biased towards stationary signals occurring over longer durations, as very short acoustic events could be smoothed out by the window size of the CNN. To achieve a similar embedding which includes information from higher frequencies and can receive variable length inputs one could retrain the CNN. However, doing so would require acquiring an even larger dataset than the 2,084,320 annotated clips (over 5000 hours) used by Google, and therefore a hybrid transfer learning approach would be more feasible.

### Dimensionality reduction

To produce Fig. 2 we used UMAP^37^ to embed the 128-dimensional acoustic features into a two-dimensional space. For the global comparison (Fig. 2a) there was a large sample size imbalance among the datasets. To ensure the dimensionality reduction was not biased, we randomly subsampled 27.6 hours of data from each dataset before running the UMAP algorithm, then all points were re-projected into 2D based on this embedding.

### Multi-class classification

We performed multi-class classification using a random forest classifier^46^ with 100 trees on acoustic features averaged over five minutes. We used a five-fold cross validation procedure in which data were split into stratified training and test sets using an 80:20 ratio. The balanced accuracy of the classifier on the test set was reported as average F1 score for each class to account for sample size imbalances among classes.

### Quantifying biodiversity and habitat quality

In Ithaca, USA between 25 February and 31 August 2017 near-continuous recordings were made using Swift recorders across 30 sites through the Sapsucker Woods area at a sample rate of 48 kHz. For each one-hour period of each day during this period, we randomly selected one out of the 30 sites in which to quantify biodiversity within the audio recording. For the chosen site and hour combination, a one hour audio clip was manually annotated to pick out all avifaunal species vocalising. Values of avian richness were normalised by sampling effort for all sites. Annotations were made using the Raven Pro software^47^.

For each of the 17 sites across a logged tropical forest biome in Sabah, Malaysia we estimated above ground biomass (AGB, log_10_(t.ha^−1^)). Raw AGB values across the landscape were taken from Pfeifer et. al.’s estimates based on ground surveys of the same study site^48^. For each of our recording locations we averaged AGB from all points within 1 km of the recorder.

### Anomaly score definition and density estimation

We used a Gaussian Mixture Model (GMM) with 10 components and diagonal covariance matrices to fit a probability density function to five days of acoustic features from each site (449,280 clips of 0.96 s per site). Acoustic features were calculated at the 0.96 s resolution with no averaging over longer time windows, in the full 128-dimensional feature space. We tested for improvements to the method by estimating the probability distribution using: (i) additional GMM components, (ii) non-diagonal covariance matrices, and (iii) using a Dirichlet-process Bayesian GMM^49^. Each of these modifications delivered only small advantages (with respect to the ability to identify synthetic anomalies) despite considerable increases in computational complexity. Accordingly, here we report the results of a 10-component GMM with diagonal covariance matrices in the 128-feature space.

The anomaly score of each 0.96 s audio clip was defined as the negative log likelihood of its acoustic feature vector, given the probability density function for the site at which the audio was recorded.

We used the GMM as a data exploration tool to pull out the most anomalous and typical sounds over a five day period in a logged forest site in Sabah, Malaysia (Figs 4a, 4b). To characterise the most typical sounds of the soundscape, we found the audio clips from the five day period which were closest (Euclidean distance) to each of the 10 GMM components in the feature space. To find a small set of distinct anomalous sounds we first clustered the 50 most anomalous audio clips using affinity propagation clustering^50^, which returns a variable number of clusters. Then, from each of the clusters we picked the clip which had the maximum anomaly score as a representative for the final list of anomalies.

In Fig. 4a, we show a 2-dimensional representation of a 128-dimensional acoustic feature space in which the GMM-derived probability density function is depicted from a logged tropical forest site in Sabah, Malaysia. Dimensionality reduction was performed by applying principal component analysis (PCA) to the five days of 0.96 s audio clips used to fit the GMM. Anomalous points and the centres and covariances of each of the GMM components was projected into 2D using the same embedding, and shaded areas represent two standard deviations from each of the centres. PCA was used over other non-linear dimensionality reduction techniques to enable straight-forward visualisation of the probability density function.

### Anomaly playback experiments

Three variants from the following five categories of sounds were used for the anomaly playback experiments; chainsaws, gunshots, lorries, chopping, talking. All sounds were played in WAV format on a Behringer Europort HPA40 Handheld PA System, and the audio files and speaker together were calibrated to the following sound pressure levels at 1m (chainsaws: 110 dB SPL, gunshots: 110 dB SPL, lorries: 90 dB SPL, chopping: 90 dB SPL, talking: 65 dB SPL). Real world SPL levels are higher for chainsaws and gunshots, but we were unable to reproduce sound pressure levels above 110 dB SPL with the speaker used. All fifteen playback sounds were played whilst holding the speaker at hip height facing an Audiomoth recording device affixed to a tree at chest height. This was repeated at distances of 1m, 10m, 25m, 50m and 100m.

## Acknowledgements

We would like to thank Till Hoffman for his input in selecting the audio features. Thanks to the field staff and organisations who enabled the data collection from all our sites: *Sabah:* Jani Sleutel, Nursyamin Zulkifli, Adi Shabrani, Dr. Henry Bernard, SAFE Project, *Ithaca:* Ray Mack, Ben Thomas, Cornell Lab of Ornithology, *Congo:* Phael Malonga, Frelcia Bambi, Elephant Listening Project/Wildlife Conservation Society, *New Zealand:* Mike Ogle, New Zealand Department of Conservation, *Sulawesi:* Indonesian Ministry of Research. Data from Sulawesi were collected under permit number: 2881/FRP/E5/Dit.KI/VII/2018. Data from the Republic of Congo were collected with permission of the Republic of Congo Ministry of Forestry.

## Authors contributions

S.S.S., N.S.J., B.D.F., L.P. and R.M.E. all contributed to the conceptualisation, development of analysis methods and final implementation of this study. S.S.S., L.P., R.M.E., D.J.C., H.K., C.D.L.O. and P.H.W. contributed to eco-acoustic data collection. S.S.S., N.S.J., C.D.L.O. and R.M.E. led the manuscript writing process, with input provided from all authors.

## Data and materials availability

Code to reproduce results and figures from this study is available on Zenodo at https://doi.org/10.5281/zenodo.3530203 and the associated data can be found at https://doi.org/10.5281/zenodo.3530206.

## Supplementary Materials

**Supplementary Audio File 1:** a short audio file containing 0.96 s clips of 12 anomalies and 10 typical sounds detected from a five day period of monitoring in Sabah, MY is attached to this manuscript as ‘12_anoms_10_typical_sounds_sabah_my.wav’.

**Supplementary Audio File 2:** a short audio file containing 0.96 s clips of four anomalies and four typical sounds highlighted in Fig. 4a is attached to this manuscript as ‘fig4a_4_anoms_4_typical_sounds_sabah_my.wav’.

**Figure S1:**
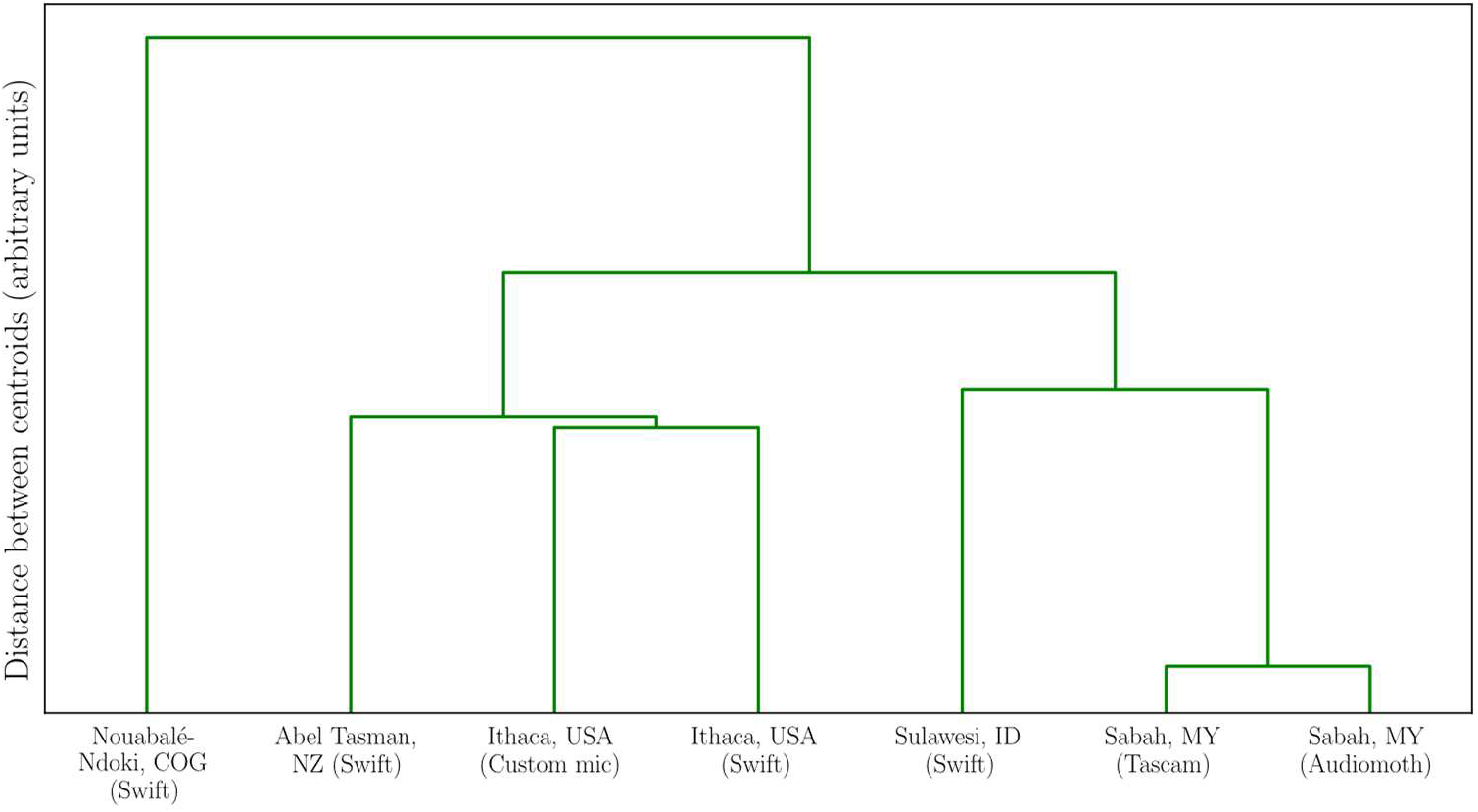
Similar biomes cluster in acoustic feature space, regardless of recording technology used. We embedded eco-acoustic data recorded using a variety of technologies from five different ecosystems in a common high-dimensional acoustic feature space and used UMAP to visualise the features in 2D (Fig 2a). We used hierarchical clustering to compare the centroids of each dataset, and found similar biomes to cluster closely when visualised as a dendrogram. Southeast Asian tropical biomes (Sabah, MY and Sulawesi, ID) formed one cluster, and temperate regions (Abel Tasman National Park, NZ and Ithaca, USA) formed another. This result was independent of recording device used.

**Figure S2:**
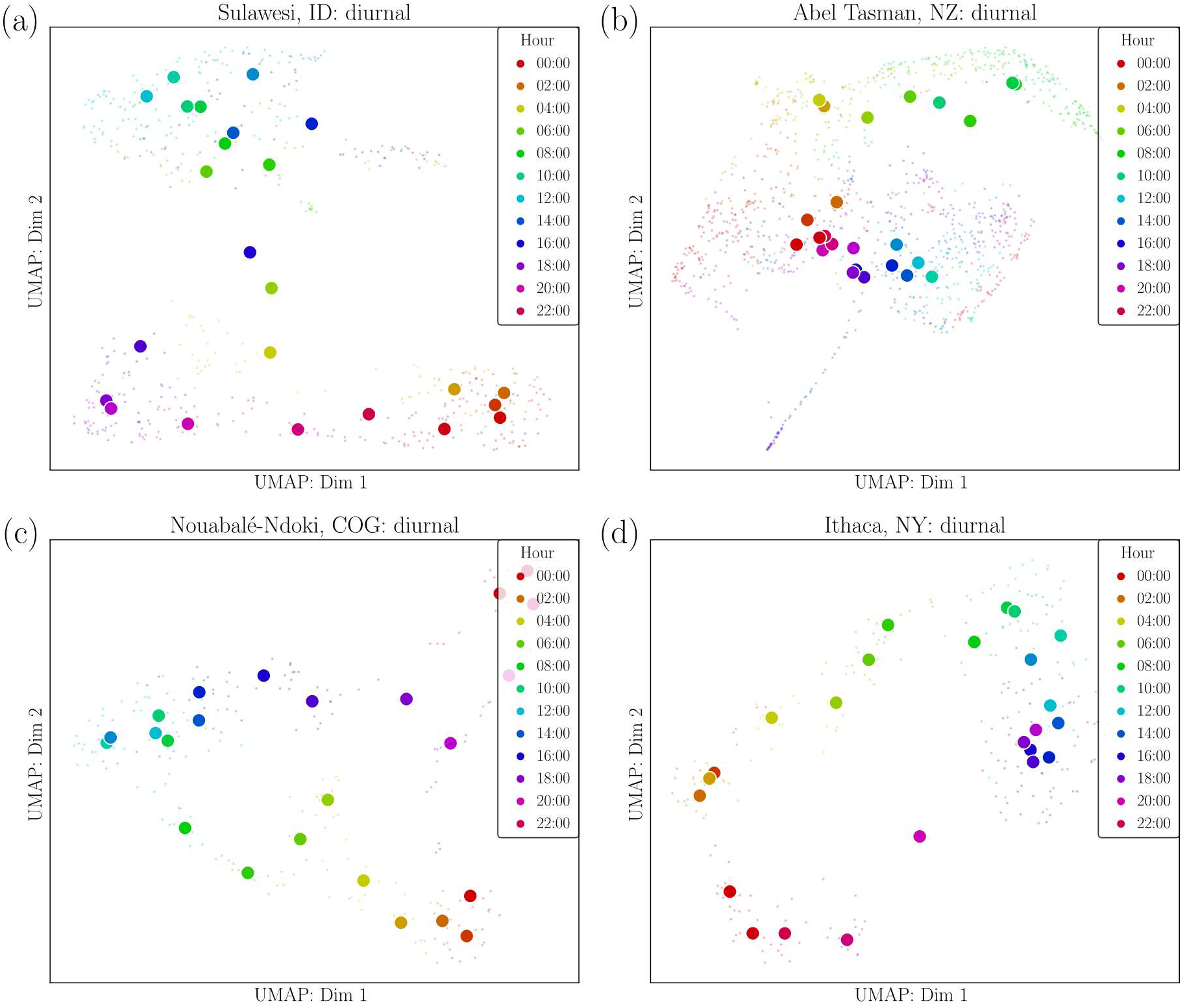
Unsupervised dimensionality reduction using a common feature-set reveals diurnal cycling patterns across ecosystems. Acoustic data from a single site within four different ecosystems was embedded in the same acoustic feature space using a pre-trained convolutional neural network: **(a)** Sulawesi, Indonesia, **(b)** Abel Tasman National Park, New Zealand, **(c)** Nouabalé-Ndoki National Park, Republic of Congo, **(d)** Ithaca, USA. This embedding shows clear diurnal cycling from each of the ecosystems, as shifts in the vocalising animals throughout the different hours of the day result in a smoothly varying acoustic signature. In all panels, uniform manifold learning technique (UMAP)^37^ was used to visualise a 2D embedding from the full 128-dimensional acoustic feature space, and centroids of classes are denoted by larger points.

**Figure S3:**
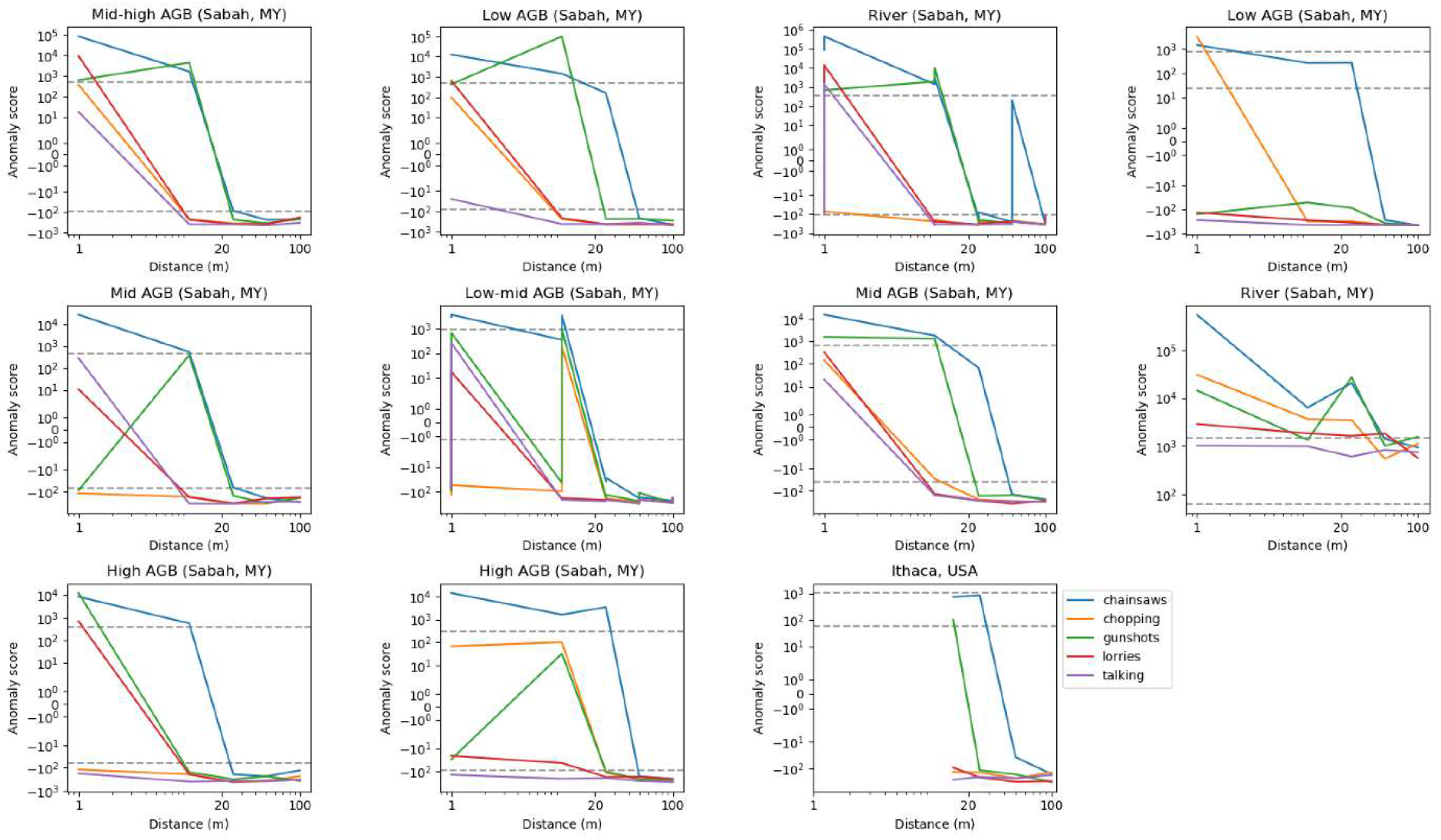
Unsupervised anomaly detection across a range of habitat qualities in Sabah, Malaysia and Ithaca, USA. We tested the sensitivity of our unsupervised anomaly detection algorithm across 10 sites from a variety of habitat qualities at a tropical forest site in Sabah, and a temperate forest in Ithaca. Largely sounds such as chainsaws and gunshots were identified as anomalous at distances of up to 25 m, however at further distances the playback sounds were barely audible over background noise (Fig. 4d). The anomaly detection routine struggled at the tropical forest site near a river, where all playback sounds were identified as anomalous. In Ithaca, due to physical constraints we could not perform the playback experiments at distances closer than 15 m from the recorder.

## References

1 Patino, M. et al. Accuracy of acoustic respiration rate monitoring in pediatric patients. Pediatr. Anesth. 23, 1166–1173 (2013).

2 Cullington, D. W., MacNeil, D., Paulson, P. & Elliott, J. Continuous acoustic monitoring of grouted post-tensioned concrete bridges. NDT E Int. 34, 95–105 (2001).

3 Harma, A., McKinney, M. F. & Skowronek, J. Automatic surveillance of the acoustic activity in our living environment. in 2005 IEEE International Conference on Multimedia and Expo 4 pp.- (2005). doi:10.1109/ICME.2005.1521503.

4 Atlas, L. E., Bernard, G. D. & Narayanan, S. B. Applications of time-frequency analysis to signals from manufacturing and machine monitoring sensors. Proc. IEEE 84, 1319–1329 (1996).

5 Vitousek, P. M. Beyond global warming: Ecology and global change. Ecology 75, 1861–1876 (1994).

6 Rapport, D. J. What constitutes ecosystem health? Perspect. Biol. Med. 33, 120–132 (1989).

7 Rapport, D. J., Costanza, R. & McMichael, A. J. Assessing ecosystem health. Trends Ecol. Evol. 13, 397–402 (1998).

8 Fitzpatrick, M. C., Preisser, E. L., Ellison, A. M. & Elkinton, J. S. Observer bias and the detection of low-density populations. Ecol. Appl. 19, 1673–1679 (2009).

9 Hampton, S. E. et al. Big data and the future of ecology. Front. Ecol. Environ. 11, 156–162 (2013).

10 Soranno, P. A. & Schimel, D. S. Macrosystems ecology: Big data, big ecology. Front. Ecol. Environ. 12, 3–3 (2014).

11 Baret, F. & Buis, S. Estimating canopy characteristics from remote sensing observations: Review of methods and associated problems. in Advances in Land Remote Sensing: System, Modeling, Inversion and Application (ed. Liang, S.) 173–201 (Springer Netherlands, 2008). doi:10.1007/978-1-4020-6450-0_7.

12 Sollmann, R., Mohamed, A., Samejima, H. & Wilting, A. Risky business or simple solution – Relative abundance indices from camera-trapping. Biol. Conserv. 159, 405–412 (2013).

13 Gemmeke, J. F. et al. Audio Set: An ontology and human-labeled dataset for audio events. Google AI https://ai.google/research/pubs/pub45857 (2017).

14 Hershey, S. et al. CNN architectures for large-scale audio classification. in 2017 IEEE International Conference on Acoustics, Speech and Signal Processing (ICASSP) 131–135 (2017). doi:10.1109/ICASSP.2017.7952132.

15 Gibb, R., Browning, E., Glover‐Kapfer, P. & Jones, K. E. Emerging opportunities and challenges for passive acoustics in ecological assessment and monitoring. Methods Ecol. Evol. 10, 169–185 (2019).

16 Bravo, C. J. C., Berríos, R. Á. & Aide, T. M. Species-specific audio detection: A comparison of three template-based detection algorithms using random forests. PeerJ Comput. Sci. 3, e113 (2017).

17 Stowell, D., Wood, M., Stylianou, Y. & Glotin, H. Bird detection in audio: A survey and a challenge. ArXiv160803417 Cs (2016).

18 Stowell, D., Benetos, E. & Gill, L. F. On-bird sound recordings: Automatic acoustic recognition of activities and contexts. ArXiv161205489 Cs (2016).

19 Towsey, M., Planitz, B., Nantes, A., Wimmer, J. & Roe, P. A toolbox for animal call recognition. Bioacoustics 21, 107–125 (2012).

20 Aide, T. M. et al. Real-time bioacoustics monitoring and automated species identification. PeerJ 1, e103 (2013).

21 Maher, R. C. Acoustical Characterization of Gunshots. in 2007 IEEE Workshop on Signal Processing Applications for Public Security and Forensics 1–5 (2007).

22 Stowell, D., Petrusková, T., Šálek, M. & Linhart, P. Automatic acoustic identification of individual animals: Improving generalisation across species and recording conditions. ArXiv181009273 Cs Eess (2018).

23 Pijanowski, B. C. et al. Soundscape ecology: The science of sound in the landscape. BioScience 61, 203–216 (2011).

24 Sueur, J., Pavoine, S., Hamerlynck, O. & Duvail, S. Rapid acoustic survey for biodiversity appraisal. PLOS ONE 3, e4065 (2008).

25 Eldridge, A. et al. Sounding out ecoacoustic metrics: Avian species richness is predicted by acoustic indices in temperate but not tropical habitats. Ecol. Indic. 95, 939–952 (2018).

26 Fuller, S., Axel, A. C., Tucker, D. & Gage, S. H. Connecting soundscape to landscape: Which acoustic index best describes landscape configuration? Ecol. Indic. 58, 207–215 (2015).

27 Sueur, J. Indices for Ecoacoustics. in Sound Analysis and Synthesis with R (ed. Sueur, J.) 479–519 (Springer International Publishing, 2018). doi:10.1007/978-3-319-77647-7_16.

28 Mammides, C., Goodale, E., Dayananda, S. K., Kang, L. & Chen, J. Do acoustic indices correlate with bird diversity? Insights from two biodiverse regions in Yunnan Province, south China. Ecol. Indic. 82, 470–477 (2017).

29 Bohnenstiehl, D., Lyon, R., Caretti, O., Ricci, S. & Eggleston, D. Investigating the utility of ecoacoustic metrics in marine soundscapes. J. Ecoacoustics 2, R1156L (2018).

30 Bradfer-Lawrence, T. et al. Guidelines for the use of acoustic indices in environmental research. Methods Ecol. Evol. 0, (2019).

31 Gavin, M. C., Solomon, J. N. & Blank, S. G. Measuring and monitoring illegal use of natural resources. Conserv. Biol. 24, 89–100 (2010).

32 Clavero, M. & García-Berthou, E. Invasive species are a leading cause of animal extinctions. Trends Ecol. Evol. 20, 110 (2005).

33 Walther, G.-R. et al. Ecological responses to recent climate change. Nature 416, 389–395 (2002).

34 Sala, O. E. et al. Global biodiversity scenarios for the year 2100. Science 287, 1770–1774 (2000).

35 Hill, A. P. et al. AudioMoth: Evaluation of a smart open acoustic device for monitoring biodiversity and the environment. Methods Ecol. Evol. 9, 1199–1211 (2018).

36 Ewers, R. M. et al. A large-scale forest fragmentation experiment: The Stability of Altered Forest Ecosystems Project. Philos. Trans. R. Soc. Lond. B Biol. Sci. 366, 3292–3302 (2011).

37 McInnes, L., Healy, J. & Melville, J. UMAP: Uniform Manifold Approximation and Projection for dimension reduction. ArXiv180203426 Cs Stat (2018).

38 Chinchor, N. MUC-4 Evaluation Metrics. in Proceedings of the 4th Conference on Message Understanding 22–29 (Association for Computational Linguistics, 1992). doi:10.3115/1072064.1072067.

39 Gunnarsson, T. G., Gill, J. A., Newton, J., Potts, P. M. & Sutherland, W. J. Seasonal matching of habitat quality and fitness in a migratory bird. Proc. R. Soc. B Biol. Sci. 272, 2319–2323 (2005).

40 Aide, T. M., Zimmerman, J. K., Herrera, L., Rosario, M. & Serrano, M. Forest recovery in abandoned tropical pastures in Puerto Rico. For. Ecol. Manag. 77, 77–86 (1995).

41 Papán, J., Jurečka, M. & Púchyová, J. WSN for forest monitoring to prevent illegal logging. in 2012 Federated Conference on Computer Science and Information Systems (FedCSIS) 809–812 (2012).

42 Hrabina, M. & Sigmund, M. Acoustical detection of gunshots. in 2015 25th International Conference Radioelektronika (RADIOELEKTRONIKA) 150–153 (2015). doi:10.1109/RADIOELEK.2015.7128993.

43 Sethi, S. S., Ewers, R. M., Jones, N. S., Orme, C. D. L. & Picinali, L. Robust, real-time and autonomous monitoring of ecosystems with an open, low-cost, networked device. Methods Ecol. Evol. (2018) doi:10.1111/2041-210X.13089.

44 Villanueva-Rivera, L. J., Pijanowski, B. C., Doucette, J. & Pekin, B. A primer of acoustic analysis for landscape ecologists. Landsc. Ecol. 26, 1233 (2011).

45 Pieretti, N., Farina, A. & Morri, D. A new methodology to infer the singing activity of an avian community: The Acoustic Complexity Index (ACI). Ecol. Indic. 11, 868–873 (2011).

46 Breiman, L. Random Forests. Mach. Learn. 45, 5–32 (2001).

47 Charif, R., Waack, A. & Strickman, L. Raven Pro 1.4 user’s manual. Cornell Lab Ornithol. Ithaca NY 25506974, (2010).

48 Pfeifer, M. et al. Deadwood biomass: an underestimated carbon stock in degraded tropical forests? Environ. Res. Lett. 10, 044019 (2015).

49 Blei, D. M. & Jordan, M. I. Variational inference for Dirichlet process mixtures. Bayesian Anal. 1, 121–143 (2006).

50 Frey, B. J. & Dueck, D. Clustering by passing messages between data points. Science 315, 972–976 (2007).

